# Structural engineering of dual-targeting antisense oligonucleotides enhances exon 44 skipping for treating Duchenne muscular dystrophy

**DOI:** 10.64898/2026.10.01.754576

**Authors:** Grant Shoffner, Florian Barthelemy, Justin Pi, Suzan Manoian, Brian T. Nguyen, Isaiah Little, Stanley F. Nelson, M. Carrie Miceli, Feng Guo

## Abstract

Antisense oligonucleotides (ASOs) inducing exon-skipping during pre-mRNA splicing offer promising treatment for diseases such as Duchenne muscular dystrophy. Recently, dual-targeting ASOs that simultaneously bind two splicing regulatory sequences have been shown to improve exon-skipping efficiency. Current ASOs are designed to interact with target RNAs solely based on Watson-Crick base pairing. Here we report the development of dual-targeting three-dimensional bridge (3D-bridge) ASOs in which tertiary interactions are engineered to increase target-binding affinity and the number of target residues occupied. We designed a 3D-bridge ASO for skipping exon 44 of dystrophin mRNA and biochemically validated its enhanced binding affinity for the target RNA over a similar ASO with the tertiary interactions disrupted. We subsequently optimized the 3D-bridge ASO by varying its ends and ribose chemical modifications using patient-derived induced directly reprogrammed myotubes and identified an LNA/DNA mixmer as the most potent. We further demonstrate enhanced exon-skipping efficacy and potency of the 3D-bridge ASO compared to previously reported ASOs in myotubes differentiated from patient-derived primary myoblasts and MyoD-reprogrammed fibroblasts harboring mutations amenable to reframing through exon 44 skipping. In conclusion, the 3D-bridge ASO technology improves the current dual-targeting design, demonstrating the utility and power of structure-based ASO design methods.

**Significance:** Exon-skipping ASOs typically rely solely on Watson-Crick base pairing to hybridize to target pre-mRNAs. We introduce a structure-based approach in ASO design by incorporating additional tertiary interactions that can strengthen and increase the specificity of these oligos. This 3D-bridge ASO technology can be used for treating Duchenne muscular dystrophy and many other diseases.

## Introduction

Duchenne muscular dystrophy (DMD) is a rare, genetic disease that affects 1 in 5,000 boys, resulting from mutation of the *DMD* gene, essential for muscle health and regeneration (1–3). In DMD loss of functional dystrophin protein leads to progressive muscle weakness typically resulting in full time wheelchair use by age 12 and premature death due to respiratory or cardiac insufficiency in their 20s. (4–6). Most *DMD* mutations are large frameshifting deletions (around 65%), followed by large duplications (around 10% of the cases) and point mutations or small changes (around 25% of cases) (7) and generally lead to complete loss of functional dystrophin protein. A milder phenotype, known as Becker muscular dystrophy (BMD), is most commonly caused by in-frame deletions leading to the production of a truncated but partially functional dystrophin protein. Based on observations of functionality of BMD dystrophin proteins, a number of dystrophin replacement strategies aimed at restoring shortened in-frame dystrophin proteins have been advanced to clinical trial and/or FDA approval, including *DMD* exon skipping and micro-dystrophin gene therapy. However, these treatments are neither curative nor applicable to all DMD patients; thus, there remains a pressing unmet need.

As many as 60% of *DMD* mutations can be reframed by single exon skipping, a therapeutic strategy wherein specific *DMD* exons are targeted for exclusion during splicing (8). By employing ASOs which sterically block specific exonic splice regulatory sequences, target *DMD* exons and their neighboring introns are removed, enabling the production of a truncated but largely functional dystrophin (9, 10). Currently there are four FDA-approved ASOs to treat DMD patients amenable to exon-skipping of 45, 51, and 53, which can treat 8%, 13%, and 8% of DMD patients, respectively (11–16). All four drugs are phosphorodiamidate morpholino oligomers (PMOs). Despite this progress, the efficacy of these ASO drugs remains limited, with between 0.3% and 6% of native protein levels recovered during clinical trials and while real word data supports efficacy, none have proven to halt progression (17–19).

Development of exon 44-skipping ASOs for DMD, which have the potential to treat about 6% of patients (8) has made great strides in recent years with potential FDA approvals on the horizon. In their landmark work in 2007, Wilton *et al.* identified the first exon 44-skipping oligo H44A (Fig. 1*A*), a 24-nt ASO based on the 2’-O-Methyl (2′-OMe) chemistry and phosphorothioate (PS) backbone that targets exon 44 bases 61-84 (20). Avidity Biosciences recently reported a 26-nt PMO binding exon 44 at the upstream residues 8-33, and conjugated this ASO to an antibody against transferrin receptor 1 to enhance cellular uptake (21), with recent reports of phase1/2 DMD exon 44-skipping trials supporting the potential approval of an Avidity exon 44 skipping drug with 25% of normal level of dystrophin protein (22). In parallel, NS Pharma proposed dual-targeting ASOs that simultaneously recognize two distal regulatory sites to induce exon 44 skipping and demonstrated that these designs have superior activity over single-site targeting controls (23). Their screen identified positions 19-33, 61-75, and 91-103, as the most active windows for generating dual-targeting ASOs. The resulting lead ASO Brogidirsen (NS-089/NCNP-02), a 24-nt PMO, underwent a phase 1/2 clinical trial, which achieved 16.63% and 24.47% of normal level of dystrophin at 40 mg/kg and 80 mg/kg, respectively (24, 25).

**Fig. 1.**
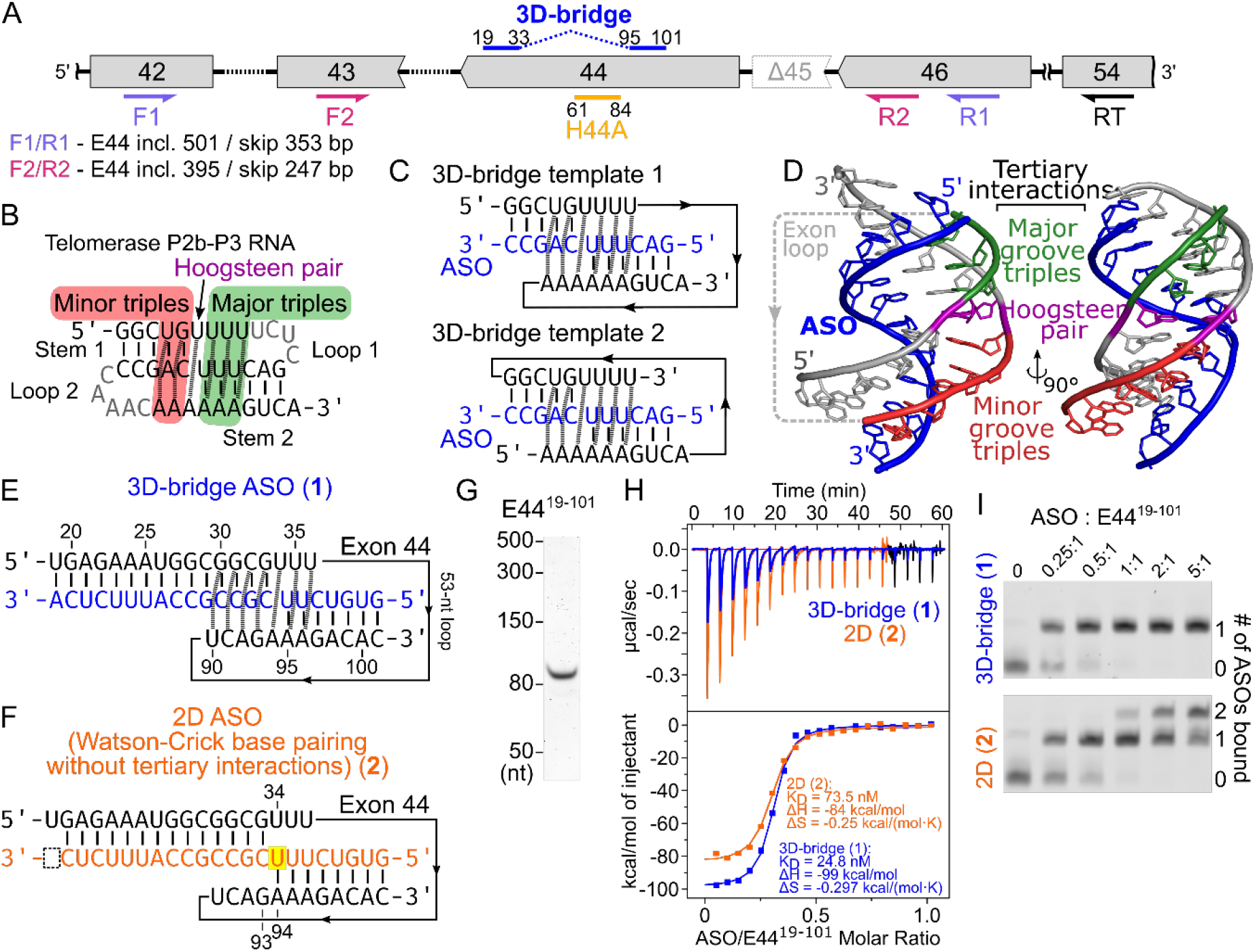
Design of 3D-bridge ASOs for dystrophin exon 44 skipping. (*A*) Region of the dystrophin pre-mRNA showing the binding sites for previously reported ASO H44A (yellow) (20), our 3D-bridge oligo (blue), reverse transcription primer (black), and nested PCR primers (pink and lavender). Exons (grey boxes) and introns (dashed lines) are not drawn to scale. Residue numberings on ASOs refer to the position within exon 44. Sizes of expected PCR products are shown. (*B*) Base-pairing (vertical bars) and tertiary (dashed lines) interactions in the telomerase RNA P2b-P3 pseudoknot (27) that inspired the 3D-bridge approach. (*C*) Altering strand connectivity of the telomerase pseudoknot yield two templates for designing ASOs (blue) that recapitulate this structure upon binding target RNA (black). (*D*) Three-dimensional model of 3D-bridge ASO (template 1) binding to target RNA, derived from the telomerase pseudoknot structure. (*E*) Design of 3D-bridge ASO targeting two ESEs of exon 44 based on template 1. (*F*) A 2D ASO that is not expected to form the tertiary interactions present in the 3D design, for comparison. (*G*) Denaturing PAGE of the purified exon 44 RNA construct (E44^19-101^) used for binding assays. Stained with SYBR-Green II. (*H*) ITC thermograms (upper panel) and isotherms (lower panel) with fitted curves for the 3D-bridge (blue) or 2D oligo (orange) titrated into E44^19-101^. (*I*) Native PAGE of E44^19-101^ annealed with ASO at varying input molar ratios.

The exciting development of dual-targeting ASOs raises new questions and opportunities for further improvement. The simultaneous binding of an ASO to two distinct target sites requires the intervening RNA sequence to loop out. The loop adjoins neighboring duplexes formed between the ASO and target RNA, creating a higher density of nucleotide residues and thereby electrostatic repulsion between their backbones and potential steric clashes. A previous effort to discover ASOs targeting flanking regions of a hairpin in the tau pre-mRNA utilized an unpaired nucleotide linker to connect the two arms of the ASO and mitigate conformational strain upon target hybridization (26). In the present study, we introduced a structural biology-based approach called three-dimensional bridge (3D-bridge) ASO to resolve this challenge for dual-targeting designs by inducing a tertiary interface between the RNA loop and ASO-RNA duplexes. We designed a 3D-bridge ASO for inducing exon 44 skipping, characterized it using biochemical assays, and optimized it in patient-derived cells. The 3D-bridge ASO displayed greater exon 44-skipping activity over previously reported ASOs.

## Results

### Concept and design templates of 3D-bridge ASO

The idea of 3D-bridge ASO was inspired by the structure of the P2b-P3 pseudoknot of human telomerase RNA (Fig. 1*B*) (27). The pseudoknot folds into a compact structure with residues in two loops engaging in two segments of major- and minor-groove base triples connected by a Hoogsteen pair. Three U residues close to the 5′ end of loop 1 contact three A-U base pairs in stem 2 in their major grooves, forming three stacked U●A-U base triples. Two A residues close to the 3′ end of loop 2 contact two Watson-Crick base pairs in stem 1 in their minor grooves. The 5′-most U residue of loop 1 hydrogen-bonds with the Hoogsteen surface of the 3′-most A residue of loop 2. This Hoogsteen pair bridges the major-groove and minor-groove triple segments. These tertiary interactions do not depend on participating residues being covalently connected as a single chain. Omitting the residues in either loop 1 or 2 (shown in gray in Fig. 1*B*) results in two separate RNA strands that can bind each other *in trans* (28).

We derived two 3D-bridge ASO design templates by assuming the central strand of the pseudoknot to be the ASO strand and the 5′- and 3′-strands to be sites on the target RNA (Fig. 1*C* and 1*D*). The target sites may be connected in two different ways, with either located upstream, thus producing design templates 1 and 2. As these design templates are generated based on an RNA structure, in principle, isosteric base triples and Hoogsteen pairs should be acceptable when designing 3D-bridge ASOs for a target RNA with a different sequence. For example, C^+^●G-C (C^+^ represents a protonated C) is a well-known major-groove base triple isosteric to U●A-U frequently observed in RNA structures (28–30).

### Design of a 3D-bridge ASO for skipping exon 44

We designed 3D-bridge ASOs targeting exon 44 that improve on previously reported oligos. Multiple ASO tiling assays of exon 44 have mapped out susceptible regions for exon skipping, including exonic splicing enhancer elements necessary for recognition by spliceosome-associated proteins (20, 23, 25). Starting from the regions 19–31/89–101 and 64–75/92–103 targeted by the most active ASOs in the NS Pharma study, we compared these sequences to our 3D-bridge templates and found that an ASO annealing to residues 19–33 and 95–101 would closely match the sequence requirements for template 1 (Fig. 1*E*). At the upstream site, a stretch of three U’s (U34–U36) provides half the needed nucleotides for forming the Hoogsteen and major-groove interactions. A corresponding string of three A’s (A94–A96) provides the second half of the required bases. Resides 89–94 should be able to participate in minor-groove base-triples with the duplex formed between ASO and RNA residues 29–33. Ideally, the intervening 52-nt exon strand (residues 37–88) is sufficiently long and unstructured to connect the major- and minor-groove strands of the structure.

We investigated the folding of exon 44 to learn if any structures in the transcript might work against binding of the 3D-bridge ASO. Predicting the secondary structure of the full 148-nt exon revealed multiple possible alternative folds, including varying long-range base pairing between the 5′ and 3′ ends. These types of long-range interactions are rarely supported by chemical probing or structure mapping approaches. Comparison of the predicted structures did show three short stem-loops between residues 48–114 that consistently occurred across all predictions (*SI Appendix*, Fig. S1*A*). To assess this prediction, we generated a fragment of the *DMD* pre-mRNA comprising exon 44 plus 25 nt of both the upstream and downstream introns. We produced the RNA by *in vitro* transcription, purified by denaturing PAGE, and performed UV melting experiments. The melting curve of exon 44 revealed three clear transitions, along with several minor features, pointing to the presence of at least three distinct structured regions (*SI Appendix*, Fig. S1*B*). We also collected the ^1^H-NOESY spectrum of a fragment of exon 44 (residues 19–101, termed E44^19-101^) at 5°C (*SI Appendix*, Fig. S1*C*). The imino-proton region revealed relatively few cross-peaks (∼12 clear peaks) that are characteristic of stable Watson-Crick or G⋅U wobble pairs. Taken together, we conclude that exon 44 likely contains several local stem-loop features but does not fold into an extended structure involving long-range pairs.

### Biochemical characterization of the 3D-bridge ASO

To determine if our engineered 3D interface enhances affinity of the 3D-bridge ASO for exon 44, we designed a second oligo with a nearly identical sequence, the same number of Watson-Crick base pairs, but disrupted tertiary interactions (Fig. 1*F*). In the “2D” oligo, we introduced an additional U residue in the ASO to base pair with A94 in the exon (highlighted base in Fig. 1*F*). This Watson-Crick base pair should sequester A94, making it unavailable for Hoogsteen pairing with U34 as in our 3D-bridge design. The neighboring G93 is not a suitable Hoogsteen partner for U34. As the Hoogsteen pair is a keystone of the P2b-P3 pseudoknot structure, preventing formation of this pair likely disrupts some or all of the adjacent base triple interactions (27, 28). To compensate for the additional U-A pair, we deleted an A at the 3’-end of the oligo that pairs with U19, so that the 3D-bridge and 2D ASOs both form ten A-U and twelve G-C Watson-Crick pairs with the exon.

We biochemically compared the association of 3D-bridge and 2D ASOs with exon 44. FWe used an exon 44 segment containing residues 19–101 (E44^19-101^) as the target RNA (Fig. 1*F*). As a proof of concept, we generated the 3D-bridge (**1**) and 2D (**2**) ASOs with RNA backbones by solid-phase synthesis and purified by reverse-phase HPLC. Isothermal titration calorimetry (ITC) binding assays showed that the 3D-bridge ASO binds E44^19-101^ with almost 3-fold tighter affinity than the 2D control (*K*_D_ = 24.8 vs 73.5 nM, Fig. 1*G* and 1*H*). This result confirms that the 3D-bridge design engages the target with higher affinity than expected for simple Watson-Crick pairing. Additionally, the 3D-bridge design binds E44^19-101^ with a larger enthalpy gain (ΔH = −99 kcal/mol vs −84 kcal/mol) and a greater entropy penalty (ΔS = −0.297 kcal/(mol·K) vs −0.25 kcal/(mol·K)) than the 2D design (Fig. 1*H*). The more favorable enthalpic gain supports the formation of engineered tertiary interactions by the 3D-bridge ASO. Commensurately, the greater entropic penalty (ΔS) reflects a substantial loss of conformational flexibility upon binding, as the 3D-bridge restricts more target residues into ordered, structurally stable states. The titration peaks for the 3D-bridge and 2D ASOs (Fig. 1*H*, top panel) are noticeably broad with delayed return to baseline, indicating relatively slow binding kinetics. This observation aligns with their dual-targeting mechanism, which requires engaging two distal sites of the target RNA to complete binding. We derived kinetic parameters from the thermograms using the AFFINImeter software (31). The association rates (*k*_on_) for the 3D-bridge and 2D oligos are comparable at 1.4 × 10^5^ M⁻^1^s⁻^1^ and 1.3 × 10^5^ M⁻^1^s⁻^1^, respectively. However, the 3D-bridge exhibits an approximate two-fold slower dissociation rate (*k*_off_, 1.4 × 10^2^ s⁻^1^) compared to the 2D ASO (3.1 × 10² s⁻¹). This slower dissociation rate further indicates that the 3D-bridge forms a more stable complex with the target RNA.

Because dual-targeting ASOs anneal to two distal target sites, it is possible that two ASOs may bind a single target transcript with each occupying one site. To address this possibility, we assessed binding stoichiometry by annealing ASO **1** or **2** with E44^19-101^ at varying molar ratios and analyzing their complexes via native gel electrophoresis (Fig. 1*I*). We found that the 3D-bridge design (**1**) strictly forms a 1:1 complex with the exon, even when a large excess of ASO is present. Thus, the binding of 3D-bridge to the two sites on the target RNA is highly cooperative. In contrast, the 2D ASO (**2**) produces a mixture of unbound exon, 1:1, and 2:1 (ASO:exon) complexes (Fig. 1*I*, lower). The formation of 2:1 complex is evidenced by the super-shift observed as the ASO:E44^19-101^ molar ratio increases above 1:1. This result clearly demonstrates that the 2D ASO does not form a fully stable complex with the exon, allowing excess ASO to compete for binding to the second site.

### 2′-O-Me 3D-bridge ASO induces exon 44 skipping in directly reprogrammed myotubes

Next, we systematically explored the compatibility of chemical modifications with our 3D-bridge design for skipping exon 44. Whereas synthetic ASOs rely on extensive backbone and sugar modifications to enhance nuclease resistance, target-binding affinity, and/or other properties, our design templates are derived from a natural occurring RNA pseudoknot fold. Consequently, it is not guaranteed that replacing one of the three strands in this native fold with a chemically modified ASO will yield the same structural configuration. We therefore synthesized 3D-bridge ASOs incorporating a uniform phosphorothioate (PS) backbone and a variety of ribose modifications and tested their exon skipping activity using a DMD patient-derived cell culture model—inducible directly reprogrammed myotubes (iDRMs) (29, 30). This system is derived from immortalized fibroblasts from a DMD donor with an exon 45 deletion, which is amenable to reading frame rescue by exon 44 skipping. Ectopic expression of the transcription factor *MyoD* from an inducible promoter drives trans-differentiation of fibroblasts to multinucleated myotube-like cells that express *DMD* mRNA and other muscle-specific transcripts. These myotube cultures were transfected with ASOs or a non-transfected control and total RNA harvested 48 hours later. To detect exon 44 skipping, we performed RT-PCR with nested primers flanking the target exon (Fig. 1*A*) and reported “% exon skipping” based on quantitating PCR fragments representing exon 44 skipped and exon 44 included/unskipped products.

We first tested whether 2′-OMe modified 3D-bridge ASOs could induce exon 44 skipping in iDRMs (Fig. 2*B*). 3D-bridge-OMe (ASO **3**) is identical to the RNA-based ASO **1** except with 2′-OMe ribose and a PS backbone. We found that ASO **3** induced exon 44 skipping to 21% ± 2% and 38% ± 5% (mean ± S.D.) at 25 and 50 nM, respectively, significantly above the background of the non-transfected control (11.7% ± 0.4%) (Fig. 2*C* and 2*D*). We examined whether extending the 5′ end of the ASO with additional bases complementary to the exon could enhance activity, since this region forms only seven Watson-Crick base pairs. A single-base extension (ASO **4**) modestly increased activity at all concentrations, whereas a two-base extension (ASO **5**) suppressed activity (Fig. 2*C* and 2*D*). This finding points to a potentially complex relationship between the flanking helical structures and ASO efficacy.

**Fig. 2.**
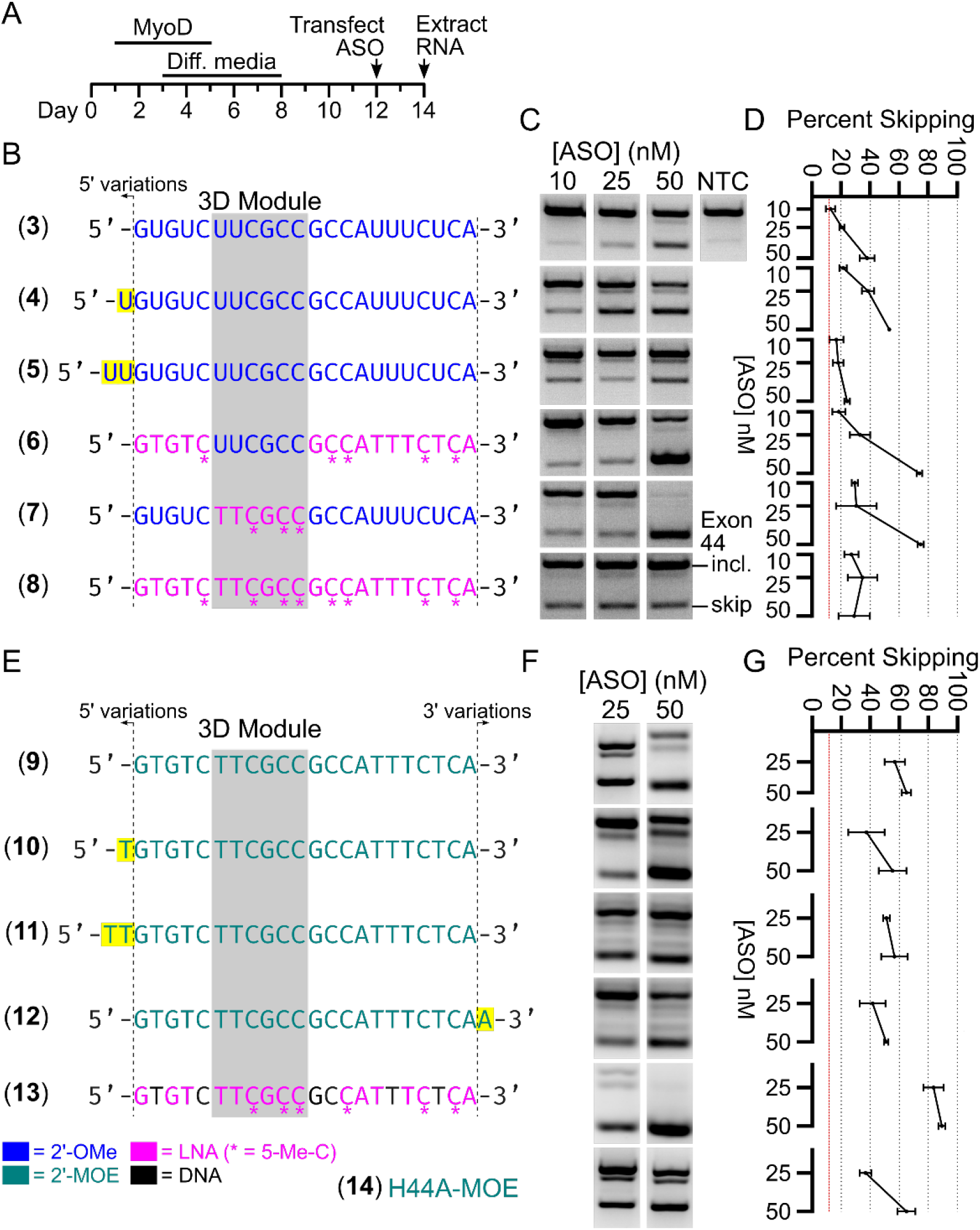
Exploration of ASO backbone chemistries in patient-derived iDRMs. (*A*) Timeline for trans-differentiation of iDRM cultures into fused multinucleated myotubes and treatment with ASO. (*B*) Panel of 3D-bridge designs testing 2′-OMe and LNA chemistries and variations in oligo length (yellow highlighted residues). (*C*) Representative agarose gel images of RT-PCR results of the dystrophin mRNA for each ASO in (*B*), with exon skipping efficiency quantified in panel (*D*). “NTC” is the non-transfected control, which sets the basal exon 44 skipping efficiency as shown by a red line in (*D*) and (*G*). (*E*) Testing 2′-MOE LNA/DNA mixmer 3D-bridge ASOs vs the previously reported ASO H44A with a 2′-MOE backbone. (*F*, *G*) Exon 44-skipping results for ASOs in (*E*). Data plotted in (*D*) and (*G*) are means ± S.D from three biological replicates.

### 3D-bridge ASO as an LNA/DNA mixmer achieves superior exon 44 skipping activity

To boost target-binding affinity without changing ASO length, we tested locked nucleic acid (LNA) and mixed LNA/2′-OMe designs. Based on the predicted tertiary structure of the ASO-RNA complex, we consider residues 6–11 of the 3D-bridge ASO to be the most essential for forming tertiary interactions with the target transcript. We term this region the “3D module” (shaded box in Fig. 2*B*), whereas the 5′ and 3′ flanking regions primarily make Watson-Crick contacts. To optimize the chemical modifications, we synthesized ASOs **6**–**8** by replacing 2′-OMe with LNA in either the flanking region, the 3D module, or both. Compared to the fully 2′-OMe and LNA ASOs **3** and **8**, LNA/2′-OMe mixmers **6** and **7**achieved substantially higher efficiency (74% ± 2% and 75 ± 2%, means ± S.D.) of exon skipping at 50 nM. This finding is consistent with the idea that different geometric restraints apply to the 3D module versus the flanking regions, and that testing combinations of chemistries can enhance structural compatibility.

The 2′-O-methoxyethyl (2′-MOE) modification is a promising ASO chemistry with demonstrated success in clinical applications (9, 31–34). We found that a 2′-MOE version of the 3D-bridge (ASO **9**) resulted in higher skipping efficiencies (65% ± 3% at 50 nM) than the fully 2’-OMe and LNA designs (ASOs **3** and **8**, 38% and 29%) (Fig. 2*E*–*G* vs 2*B*–*D*). Because the 2’-MOE 3D-bridge exhibited similar exon 44 skipping efficiencies (57% and 65%) at 25 and 50 nM, its activity appears to have plateaued. Thus, the LNA/2′-OMe mixmers (ASOs **6** and **7**) remained the most efficacious, achieving higher maximum skipping. We subsequently tested whether variations in the 5′ (ASOs **10**, **11**) or 3′ end (ASO **12**) of the 2′-MOE 3D-bridge could improve activity, and observed moderate decreases compared to ASO **9** (Fig. 2*E–G*). Notably, the 2′-MOE 3D-bridge ASO exhibited higher activity than a 2′-MOE version of the previously reported H44A (H44A-MOE, ASO **14**; the original H44A used 2′-OMe residues) (20) at 25 nM (57% vs 37%), indicative of a higher potency.

We investigated whether a more conventional LNA/DNA mixmer might offer further improvement. Both LNA/2′-OMe mixmer 3D-bridge ASOs **6** and **7** achieved the highest activity so far (∼75%). However, compared to the all-2′-OMe ASO **3,** ASO **6** introduced LNA modifications only in the six positions in the 3D module, whereas ASO **7** used 16 LNA residues in the flanking regions. Therefore, we generated LNA/DNA mixmer ASO **13** by retaining LNA at positions 6–11 and interspersing LNA and DNA in the flanking regions. Remarkably, this 3D-bridge-LNA/DNA ASO demonstrated superior exon 44 skipping activity, achieving 89% ± 2% skipping at 50 nM. The fact that ASO **13** dramatically outperforms the all-LNA ASO **8** suggests that the flexibility afforded by the DNA residues may be crucial for proper folding of the ASO-target RNA complex.

As our preliminary experiments found close to saturating activity for the 3D-bridge-LNA/DNA ASO at 25 and 50 nM, we next assessed lower concentrations to better gauge the potency of the oligo in this assay (Fig. 3). As a benchmark, we tested the 3D-bridge (**13**) against the H44A sequence with 2’-OMe modified sugars and a phosphorothioate backbone (oligo **17**), as originally reported. We transfected ASOs in quadruplicate into iDRM-based myotubes and extracted RNA 48 hours later for RT-PCR analysis. We found that the 3D-bridge-LNA/DNA (**13**) showed clear activity at 2 nM (lowest concentration tested, Fig. 3*A*). Quantification of the bands enabled us to estimate the EC_50_ of the oligo to be 2 ± 2 nM (mean ± S.D., Fig. 3*B*). While H44A achieved appreciable skipping activity at 25 and 50 nM, we were unable to confidently fit the response with the EC_50_ model. Despite this, the results demonstrate an order-of-magnitude improvement in exon 44 skipping activity for the 3D-bridge ASO relative to H44A in the iDRM myotube system.

**Fig. 3.**
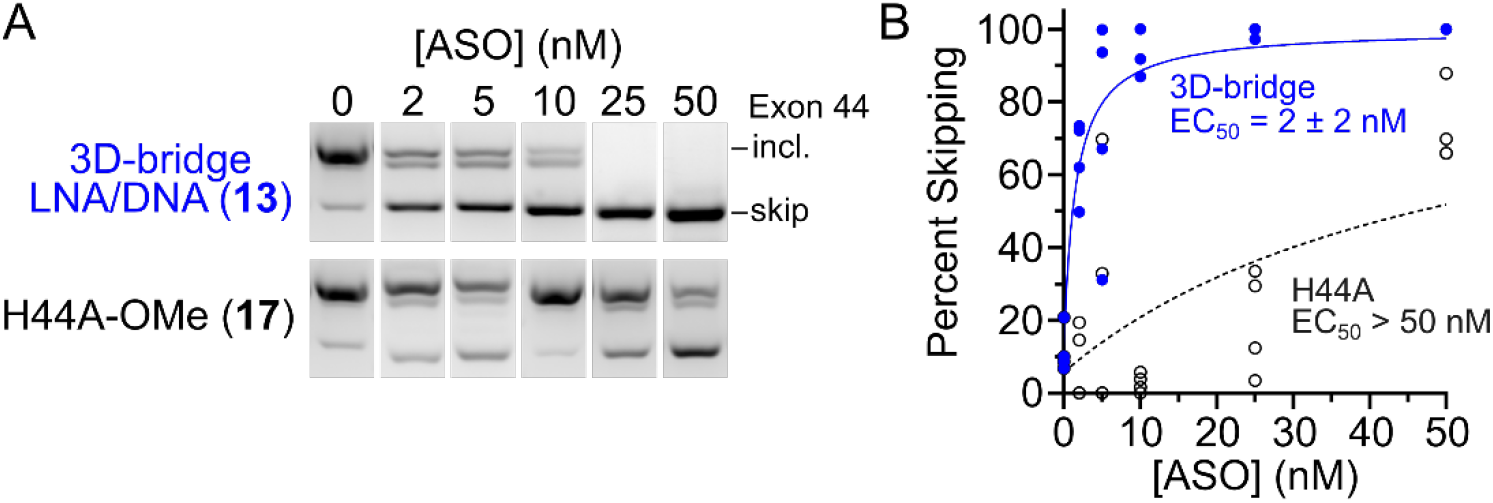
Estimate of 3D-bridge potency versus the H44A benchmark in iDRM cultures. (*A*) Representative RT-PCR gel slices from four replicate transfections of the 3D-bridge LNA/DNA mixmer (**13**, top row) or H44A-OMe ASO (**17**) in late iDRMs. (*B*) Quantification of the results in (*A*) plotted as blue dots (3D-bridge) or black circles (H44A). EC_50_ values were estimated by least-squares fitting against the data (blue and black curves). The 3D-bridge EC_50_ is reported as mean ± S.D. (n = 4). As the response for H44A was not well fit by the model, only a lower bound on EC_50_ is shown.

### 3D-bridge-LNA/DNA outperforms the 2D ASO in DMD primary myoblasts

Based on the initial success of the 3D-bridge designs in iDRM cultures, we aimed to replicate these results in alternate DMD models. We used cultured primary myoblasts from a DMD donor (patient 7022) within differentiation media to induce fusion and myotube formation (Fig. 4*A*). We first directly compared 3D-bridge-LNA/DNA (ASO **13**) with a scrambled control (ASO **15**) (Table 1). As expected, increased concentration of the scrambled control, does not lead to increased exon-skipping; only the baseline level of endogenously skipped transcript is observed. Conversely, 3D-bridge-LNA/DNA, reaches a plateau at approximately 80% skipping at 25 nM. Since the initial concentrations resulted in early saturation, we subsequently lowered the concentration to more accurately assess ASO potency. We tested and compared the 3D-bridge ASO **13** to the 2D ASO **16**. The 2D ASO **16** is identical in sequence to ASO **2** but features an LNA/DNA mixmer chemistry closely matching that of ASO **13**. These ASOs were tested in two patient-derived primary myoblasts (patient 7022 and 7021), both harboring an exon 45 deletion, but otherwise represent distinct DMD breakpoints and genetic backgrounds.

**Fig. 4.**
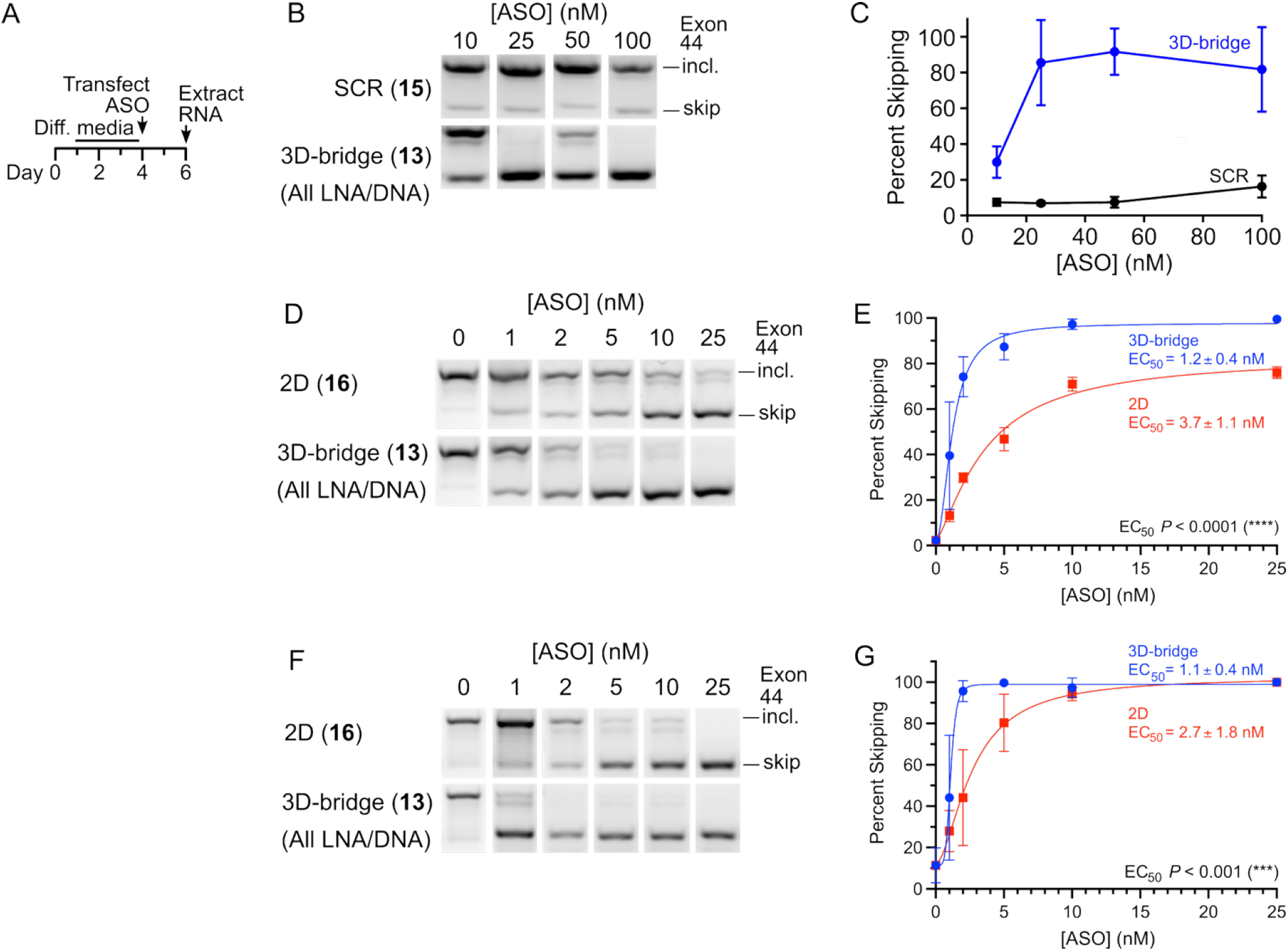
Estimation of potency of 3D-bridge oligo verses the 2D in primary myoblast cultures from two different patients. (*A*) Experimental timeline for differentiation and transfection. (*B-E*) Exon-skipping results in patient 7022. (*B*) Representative RT-PCR gel slices from three replicate transfections. (*C*) Quantification of the results in (*B*) plotted as blue dots (3D-bridge) or black squares (scramble), (*F, G*) Exon-skipping results for patient 7021. Data plotted in (*E*) and (*G*) are reported as means ± S.D. (n = 4), where EC_50_ values were estimated by least-squares fitting against the data (blue and red curves). EC_50_ values were compared using the extra sum-of-squares F-test, where *P* <0.0001, and *P* <0.001, respectively. (*D*) and (*F*) are representative RT-PCR gel slices from four replicate transfections.

**Table 1.**
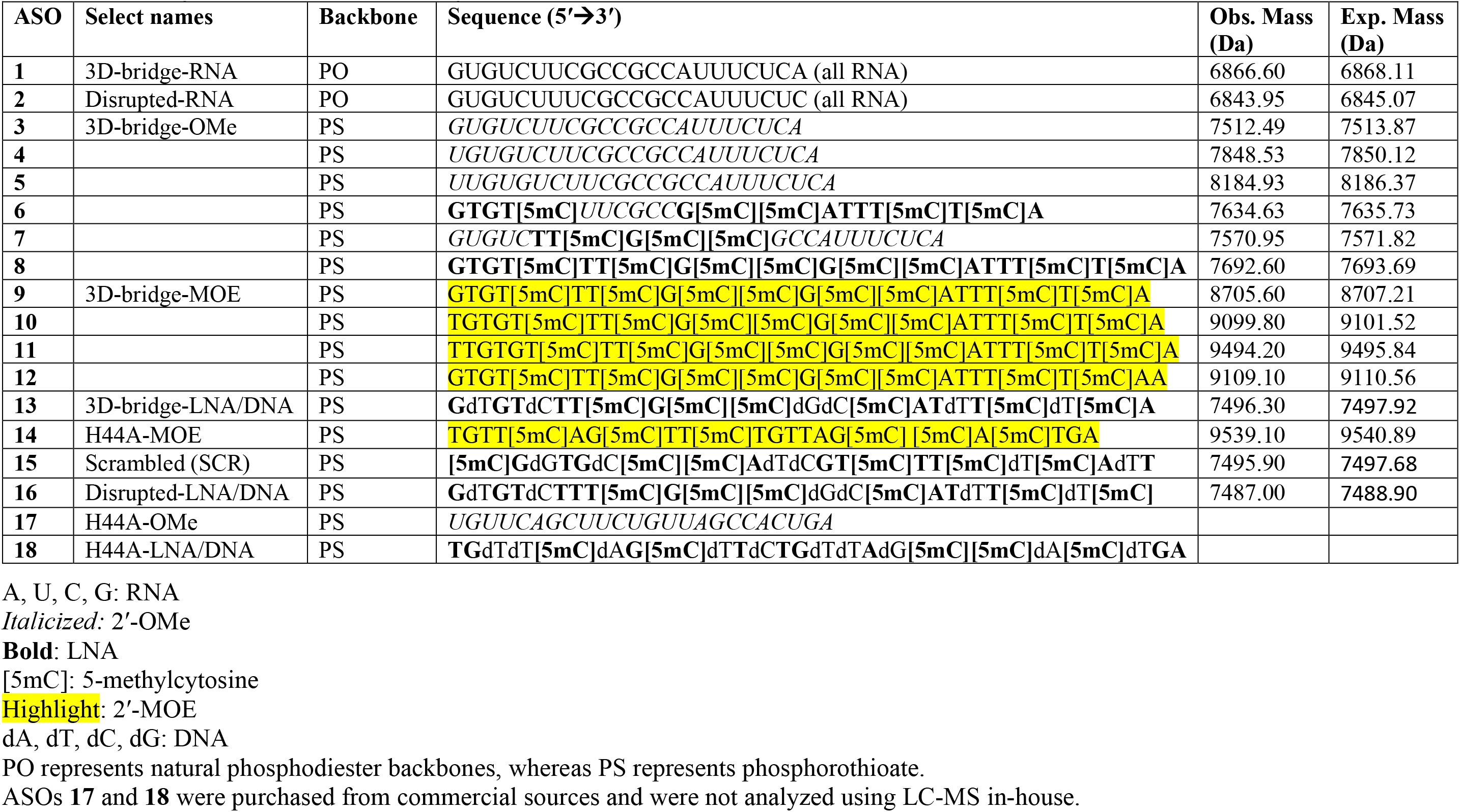
Summary of ASOs used in the study.

EC_50_ values of the 2D ASO in patients 7022 and 7021, respectively, were determined to be 3.7 ± 1.1 nM, and 2.7 ± 1.8 nM (Fig. *5C* and *5E*). Previous characterization of these patient-derived cells showed that patient 7021 has higher levels of endogenously skipped mRNA (51), which is also recapitulated in this experiment. This observation could explain why the 2D ASO in 7021 achieved near-complete skipping at 10 nM, whereas the 2D ASO in patient 7022 was slightly less potent, reaching only about 80 percent skipping. Conversely, the EC_50_ values for the 3D-bridge ASO are 1.2 ± 0.4 nM, and 1.1 ± 0.4 nM (Fig. *5C* and *5E*), which is consistent with our experiment in the iDRMs (Fig. *3*). Regardless of the higher endogenous-skipping levels in patient 7021, the 3D-bridge oligo reaches plateau in both patients at around 5 nM. Additionally, the 3D-bridge ASO was approximately 3-fold more potent than the 2D oligo in both patients, similar to the 3-fold increase in binding affinity observed in our binding assays. Finally, we performed an extra sum-of-squares F test to compare the EC_50_ values of patient 7022 and 7021, and determined the *P* values to be <0.0001 and <0.001, respectively. Together, these results demonstrate that the engineered contacts of the 3D-bridge increase the potency of an ASO relying only on Watson-crick base pair interactions by about 3-fold in a primary myoblast model.

## Discussion

In this study, we demonstrate that designing ASOs in three-dimensional space offers significant advantages over methods relying solely on Watson-Crick base pairing, especially in the context of dual-targeting strategies. Our 3D-bridge ASO binds target RNA with higher affinity (Fig. 1*H*), and its two arms engage their corresponding hybridization sites with strong cooperativity (Fig. 1*I*). These features help overcome the intrinsic limitations of conventional ASO designs. With only Watson-Crick base pairing, target-binding specificity is inherently limited because every nucleotide added to the ASO to enhance target affinity simultaneously increases the likelihood of hybridization to off-target RNAs. This off-target binding is a primary driver of toxicity and thus imposes a significant constraint on the ceiling of the therapeutic window. By engineering tertiary interactions that drive the folding of ASO-target complexes into higher-order structures, our 3D-bridge design allows the ASO to recognize additional residues. This confers a greater capacity to discriminate between on- and off-targets, providing a biophysical basis for an expanded therapeutic window. Furthermore, achieving therapeutic effects with dual-targeting ASOs requires simultaneous binding to both target sites, making hybridization to the weaker site a limiting factor for potency. The conventional solution is to add ASO nucleotides to increase base pairing with the weaker site. This approach lengthens the drug candidates, narrows therapeutic window, and increases the manufacturing costs. In contrast, the enhanced binding affinity and cooperativity of 3D-bridge ASOs boost potency with fewer residues and circumvent the length penalty typical of dual-targeting designs. For example, in their pioneering study, Watanabe *et al.* selected the 24-mer NS-089 over a 26-mer candidate for further development, specifically to favor a shorter sequence (23). We project that our 22-mer 3D-bridge ASO can engage and sterically block up to eight additional residues (Fig. 1*E*), which would otherwise require a 30-mer ASO to achieve using conventional designs. In this regard, we estimate that the 3D-bridge technology may reduce the ASO lengths in general by up to 27%. Finally, the 3D-bridge ASO dissociates from the target RNA with a slower rate, which translates to an extended target residence time—a critical parameter for sustaining therapeutic effects.

The performance of 3D-bridge ASOs with different chemical modifications may be understood in the context of the telomerase pseudoknot structure (27). PS and chemical modifications at the 2′ OH are commonly used in oligonucleotides to increase stability and nuclease resistance and enhance affinity to the target RNA (32–34). 3D-bridge ASOs with 2′-OMe, 2′-MOE, and LNA modifications all show exon-skipping activities to various degrees in iDRMs (Fig. 2). This observation aligns with the fact that, in the telomerase RNA pseudoknot structure, the backbone of the strand corresponding to 3D-bridge ASO is fully exposed to solvents and is not close to other strands. To confirm this, we built a 3D model of 3D-bridge-LNA/DNA bound to exon 44 based on the pseudoknot structure (Fig. S3). Indeed, the backbone modifications are well tolerated in the model. Our investigation indicates that LNA works better than 2′-OMe in the 3D module. We think that this result may be understood by another important structural feature of 3D-bridge designs— the ASO strand participates primarily in Watson-Crick base pairing within the target RNA, whereas the target RNA itself contributes to the tertiary interactions. As such, a 3D-bridge ASO may encourage tertiary interactions by stabilizing the Watson-Crick base pairs. Among the chemical modifications tested, LNA has the highest base pairing affinity per nucleotide due to its methylene bridge that covalently connects the 2′-oxygen to the 4′-carbon of the ribose ring and locks the ribose in the C3′-endo conformation. Previous studies showed that LNA modifications can also enhance interactions in duplex structures forming triplex interactions (35–39). Nevertheless, LNA’s high affinity can potentially cause off-target effects and toxicity, hence the interspersion of DNA residues as demonstrated in the flanking regions of ASO **13** may be critical to practical designs (40, 41).

Although PMOs represent the most successful class of exon-skipping therapies to date, novel alternative chemistries are in active clinical development (42). In particular, Wave Life Sciences is advancing an exon 53 skipping lead (WVE-N531) with a stereo-chemically defined phosphoryl guanidine backbone (43), and Daiichi Sankyo is exploring a 2′-OMe/2′-O,4′-C-ethylene-bridged nucleic acid (ENA) phosphorothioate mixmer (Renadirsen) against exon 45 (44). The later example has demonstrated safety and tolerability in a mouse model (45) and is relevant to our lead 3D-bridge design as the ENA modification resembles the methylene bridge incorporated in the LNA residues of our mixmer. Similarly, Vico Therapeutics reported promising pre-clinical data on a 2′-OMe/LNA mixmer targeting exon 51 that was well-tolerated in mice (46). Our study showed that the 3D-bridge ASOs effectively induce exon 44 skipping in myotube-like cultures derived from two different patients, one patient-derived iDRM and two patient-derived primary myoblast with exon 45 deletion DNA mutations. Exon 45 deletion is the most common mutation amenable to exon 44 skipping. Rarer DMD deletions ending at exon 43, including deletions of exons 14–43, 19–43, 30–43, 35–43, 36–43, 40–43, and 42–43, are also amenable to exon 44 skipping (25, 47). In theory, these different genetic mutations could affect the folding of the *DMD* pre-mRNA and its complex with 3D-bridge ASOs. However, because the 3D-bridge target sites are less than 60 nt apart in exon 44, we expect that the 3D-bridge ASO should perform well on most if not all relevant genetic mutations. Because patients amenable to exon skipping 44 with the exon 45 deletion can exhibit higher levels of endogenous skipping (48), mutations that do not predispose to endogenous skipping may require optimized oligos afforded by increased 3D ASO activity to reach maximal activity. In addition to the potential to improve deliver and efficacy, more potent *DMD* exon skipping drugs could reduce required frequency of injections, costs and adverse events; all significant improvements for DMD treatment. Thus, further optimization of *DMD* targeted ASOs will continue to drive advances in DMD treatment. As nearly all exons contain multiple enhancer elements, we anticipate that the ability to effectively mask multiple sites with a 3D-bridge ASO will be broadly useful for improved exon skipping applications.

*In silico* modeling of 3D-bridge ASO in complex with exon 44 provides insight into how design templates 1 and 2 may be applied to simultaneously engage two target sites (Figs. 1C and S3). These templates diverge in the spatial connectivity of the two target sites. In template 1, the upstream (5′) site ends with residues that insert into the major groove of the heteroduplex formed by the 3D-bridge ASO and the downstream target sequence; whereas the downstream site starts with residues that insert into the minor groove of the heteroduplex formed by the ASO and the upstream target sequence. In the model, the distance between these ends is 32 Å, dictating the requisite spatial span for the intervening sequence. Notably, this distance depends on the number of major- and minor-groove base triples engineered. In template 2, the upstream target site ends with residues that form Watson-Crick base pairs with the 5′ end of the 3D-bridge ASO and the downstream target site starts with residues that base pair with the 3′ end of the ASO. Consequently, the spatial gap that the intervening sequence needs to expand is approximately 72 Å. These distinct parameters suggest that template 1 imposes less constraint on the length of the intervening sequence and thus is suitable for targeting sites that are either close or far apart in sequence, whereas template 2 is applicable for targeting two distal sites.

This study represents a broader initiative to apply structural biology principles to ASO design. We previously reported the development of 3D-ASOs (28), a technology similarly derived from the telomerase pseudoknot structure but wherein a different RNA segment was abstracted as the ASO strand. In the absence of engineered chemical modifications, the 3D-modules of 3D-bridge ASO and 3D-ASO, when in complex with their respective RNA targets, adopt isosteric structures. Furthermore, our 3D-ASO study provided biochemical evidence demonstrating that non-canonical base combinations beyond U●A-U and C+●G-C triples can assemble into thermodynamically stable, isosteric base triples in the context of ASO development (28). These observations align with established RNA folding precedents (30, 49) and molecular dynamic simulations (50). These findings substantially expand the applicable target space of both 3D-ASO and 3D-bridge ASO technologies. Despite these shared structural foundations, the two technologies diverge in their mechanistic execution. 3D-Bridge ASO targets sites projected to lack defined structures, and the hybridization induces the formation of tertiary interactions and higher-order structures. Conversely, 3D-ASO engages and stabilizes a pre-folded target duplex structure alongside its flanking sequences. Thus, 3D-bridge ASO and 3D-ASO are complementary to each other. Together, they establish a technological platform that enhances ASO therapeutic development by engineering tertiary interactions.

## Materials and Methods

### Production of E44^19-101^ RNA by in vitro transcription

A single-stranded DNA Milligan template for transcription of E44^19-101^ was designed by appending the T7 promoter sequence (5′-TAATACGACTCACTATAGG-3′) to exon 44 residues 19-101 and taking the reverse complement. Note that this appends two non-natural G residues to the 5′ end of the RNA to increase transcription yield. The transcription template was produced on a K&A H-8-SE nucleic acid synthesizer via standard phosphoramidite chemistry (51). The first two residues at the 5′ end of the template were synthesized with 2′-O-methyl-modified sugars to suppress untemplated nucleotide addition during run-off transcription (52). DNA phosphoramidites with standard protecting groups were purchased from Glen Research (Cat. No.: Bz-dA: 10-1000, Bz-dC: 10-1010, iBu-dG: 10-1020, and dT: 10-1030) and 2′-OMe amidites were obtained from Thermo Fisher Scientific (Cat. No. 2′-OMe-U: 24-2044-04 and 2′-OMe-iBu-G: 27-2043-04). All amidites were resuspended in anhydrous acetonitrile at 0.1 M concentration immediately before use. The template was synthesized at the 0.2 μmol scale from Glen UnySupport 1000 CPG resin (Cat. No. 20-5241-42) using the default DNA synthesis program (K&A Labs) with the terminal dimethoxytrityl (DMT) group retained for purification. Ancillary synthesis reagents included anhydrous acetonitrile (Fisher Scientific Cat. No. AA42311K2); 0.25 M 5-Benzylthio-1H-tetrazole in acetonitrile as activator (Glen Research Cat. No. 30-3170); 10% acetic anhydride in tetrahydrofuran (Cap A, Glen Research Cat. No. 40-4012); 10% pyridine and 10% N-methylimidazole in tetrahydrofuran (Cap B, Glen Research Cat. No. 40-4122); 20 mM iodine in 2% pyridine, 10% water, and 88% tetrahydrofuran (Oxidizer, Glen Research Cat. No. 40-4130); and 3% trichloroacetic acid in dichloromethane (Deblock, Fisher Scientific Cat. No. A322-500 and D143-4).

The template was cleaved from the support and deprotected in a 50% solution of aqueous methylamine in ammonium hydroxide for 20 min at room temperature followed by incubation at 65°C for 20 min. The oligo was then purified over a Glen-Pak SPE cartridge (Glen Research Cat. No. 60-5100) following the manufacturer’s directions. The eluted oligo was dried under vacuum, resuspended in water, and buffer exchanged into 10 mM Tris pH 7.5 by repeated concentration and washing in an Amicon 0.5-mL centrifugal filter device (3 kDa MWCO). Concentration was determined by UV absorbance at 260 nm, resulting in a final yield of approximately 60 nmol template. Prior to use, the template was annealed with the promoter strand (5’-TAATACGACTCACTATAGG-3’, purchased from Integrated DNA Technologies) in a reaction comprised of 50 μM template, 50 μM promoter strand, 0.1 M NaCl, 10 mM HEPES buffer pH 7.0, and 1 mM EDTA in a final volume of 200 μL. This solution was briefly heated to 95°C and then slowly cooled to room temperature to generate the promoter duplex.

For the *in vitro* transcription reaction, the 200-μL annealed template was added to a 10-mL reaction containing 3 mM NTP mix, 2.2 mg/L T7 RNA polymerase and 1× transcription buffer (40 mM Tris pH 8.0, 25 mM MgCl_2_, 4 mM dithiothreitol, and 2 mM spermidine final concentrations). The transcribed RNA was then ethanol-precipitated and purified over 10% polyacrylamide denaturing slab gel for ∼7 hours. The RNA band was visualized by UV shadowing, excised, crushed, and extracted overnight at 4°C in TEN buffer (100 mM NaCl, 10mM Tris pH 7.5, 1 mM EDTA). The extracted RNA was concentrated, then buffer-exchanged into 1× ITC buffer (20 mM Na-HEPES, 100 mM KCl, pH 7.0). Final purity was confirmed by denaturing gel stained with SYBR-Green II (Fig. 1G).

### Synthesis of antisense oligonucleotides

All ASOs were synthesized following the standard synthesis scheme outlined above. RNA, 2′-OMe, and 2′-MOE phosphoramidites were purchased from Thermo Fisher, whereas the locked nucleic acid (LNA) phosphoramidites were acquired from Glen Research. For ASOs containing deoxy-cytidine positions, we utilized acetyl protecting chemistry (Ac-dC, Glen Research Cat. No. 10-1015) on the primary amine to minimize modification of this position during deprotection (53). Oligonucleotides were cleaved and deprotected as described, and RNAs were further treated with triethylamine trihydrofluoride (TEA·3HF) for 2.5 hours at 65°C to desilylate the 2′-OH. Samples were then purified using the Glen-Pak DNA purification cartridge (Glen Research).

If needed, samples were further purified using reverse-phase chromatography with an Agilent 1200 HPLC system and semi-preparatory TARGA C18 column (Higgins Analytical) to remove incomplete cleavage products. Finally, the purity and molecular mass of the samples were confirmed using LC-MS (Agilent Quadrupole system) (*SI Appendix,* Fig. S2 and Table 1).

### Isothermal titration calorimetry

Both target RNA and ASO were prepared in 1× binding buffer consisting of 20 mM sodium HEPES pH 7.0 and 100 mM KCl. Samples were fully buffer-exchanged into 1× ITC buffer at least 3 times using centrifugal concentrators. Samples were analyzed using a Microcal iTC-200 isothermal titration calorimeter (Malvern Panalytical). ASOs at 25–50 µM were titrated into 200 µL of target RNA at 5–10 µM over 19 injections with 2 µL each. Data were analyzed and fitted using the Origin software (version 7.0, Malvern Panalytical). Fitting was achieved until the chi-square value no longer decreased. Thermodynamic parameters, such as association constant (*K*_a_) were obtained from the software. The dissociation constant (*K*_d_) was calculated using the equation: *K*_d_ = 1/*K*_a_.

To obtain the kinetic association and dissociation rates *k*_on_ and *k*_off_, raw ITC files were imported into the AFFINImeter software where data were fitted using the kinITC function with autofitting (31).

### Native gel analysis

Target RNA at 50 nM concentration was incubated with varying molar ratios of ASO 1× binding buffer on ice for one hour. Two microliters of native gel loading dye (20 mM Tris, pH 7.5, 200 mM NaCl, 6% Ficoll, 0.01% Bromophenol blue) was added to each 10-µL sample. The samples were loaded onto a prechilled mini-gel apparatus (BIO-RAD) surrounded by ice. The native 10% polyacrylamide gel was run in 1× TBE buffer at 110 V for 70 minutes. It was then incubated in 1× SYBR Green II RNA Gel Stain (Thermo Fisher Scientific) for 20 minutes and was visualized using a Typhoon FLA 9500 scanner (GE Healthcare).

### Patient information

Subject 1015/7022 lost ambulation at age 15 and harbors a deletion of *DMD* exon 45, has low levels of self-corrected exon 44 “skipped” mRNA in derived iDRM (48), and an increased frequency of clusters of dystrophin-positive revertant fibers in his skeletal muscle (54). His skin punch was performed at the age of 8, and a muscle biopsy was performed at age 15. Subject 7021 also has a deletion of *DMD* exon 45 and was still ambulatory at the time of the biopsy when he was 15 years old. High levels of self-corrected exon 44 “skipped” mRNA were observed in his derived iDRM (48).

### Collection, isolation, and reprogramming of primary cells

Skin punches and muscle biopsies were obtained with informed consent from patients of the Center for Duchenne Muscular Dystrophy (CDMD) at UCLA under University of California Los Angeles IRB-approved protocols #11-001087 and #18-001547. Needle skeletal muscle biopsies were performed and processed as previously described (55). (56). Fibroblasts were subsequently transduced with lentiviruses containing the hTERT and a tamoxifen-inducible MyoD to allow commitment to skeletal muscle lineage as previously described (56), referred to as inducible reprogrammed myotubes (iDRM) in this manuscript.

Primary myoblasts were derived from fresh biopsy muscle and were first cultured in a muscle biopsy medium (Nutrient Mixture F-10 HAM) (Sigma-Aldrich, Saint Louis, Missouri) containing 20% FBS, 1% pen/strep, and 5 ng/mL of bFGF for 2-3 weeks. After which they were cultured in Skeletal Muscle Cell Growth Medium (Promocell, Heidelberg, Germany), 20% FBS, 1% pen/strep, and 5ng/mL of bFGF.

### Differentiation and transfection of cells

iDRMs were seeded at 150,000 cells per well of 6 well plates in fibroblast growth media pre-coated for 1 hour with 0.1% porcine gelatin (Sigma, Saint Louis, MO) and MyoD expression and differentiation as previously described (57) for 5-12 days weeks before ASO transfection.

Primary myogenic derived from muscle biopsies were pre-plated for 1 hour in growth media to remove fibroblasts which adhered to the flask in the first hour and the nonadherent fraction, enriched in myoblasts, were seeded at 150,000 cells per well in skeletal muscle cells growth media in 6-well plates pre-coated for 1 hour with 0.1% porcine gelatin.

After 2 days in skeletal muscle growth media (Promocell, Heidelberg, Germany), cell reach approximately 80% conlfuence, and were induced to fusion by switching tp Skeletal Muscle Cell Differentiation Medium (Promocell, Heidelberg, Germany) for 4 days prior to returning cells to skeletal muscle growth media and transfected on day 5 with ASO.

### Antisense transfection

After differentiation, cells were transfected with ASOs using Oligofectamine transfection reagent (Thermo Fisher Scientific) according to our previously published protocol (56) and harvested 48 hours later by lysing directly in the wells with TRIzol (Thermo Fisher Scientific) and frozen for subsequent RNA isolation and exon skipping analysis.

### RNA Isolation, PCR, and quantification

Total RNA extraction was performed using the Purelink RNA mini kit (Thermo Fisher Scientific). Exon skipping analysis was performed according to our previously published protocol (56) and skipping efficiency is calculated as the ratio of skipped mRNA transcript over the total mRNA transcripts (skipped plus unskipped).

### Structural modeling of 3D-bridge-LNA/DNA bound to exon 44

To computationally model the 3D module, the telomerase pseudoknot structure (PDB: 1YMO) was used as the foundational template. Extraneous residues were first truncated using ChimeraX (58) and the remaining sequence was modified using X3DNA-DSSR (59) to match the 3D-bridge ASO and exon 44 detailed in Fig. 1E. To recapitulate the full ASO and target sequences, an A-form RNA double helix was generated using ChimeraX and appended to the 3D module on the side with minor-groove base triples. Minor-groove base triples beyond the two layers present in the template were incorporated into the model by manually positioning the corresponding residues according to validated geometry curated in the RNA Base Triple Database (49). Subsequently, modXNA (60) interfaced with Amber (61) was deployed to introduce the requisite chemical modifications to the ASO backbone, including LNA and DNA residues and PS backbone. The resulting model was subjected to 5,000 steps of energy minimization in GROMACS (62) using the steepest descent algorithm to resolve steric clashes and optimize local geometries. The intervening sequence between the two target segments in exon 44 is not modeled.

## Acknowledgments

This work was supported by grants awarded to M.C.M. and F.G., encompassing an Idea Development Award from the Department of Defense (HT9425-24-1-0628), a National Institutes of Health (NIH) R21 award (R21HD115071), and an Appel Seed Grant Award (established in honor of Josh Bohbot) administered by the Center for Duchenne Muscular Dystrophy at the University of California, Los Angeles (UCLA). F.G. independently acknowledges funding from the NIH National Institute of Allergy and Infectious Diseases (R01AI163216). G.S. and J.P. acknowledge support from the NIH National Institute of Arthritis and Musculoskeletal and Skin Diseases (NIAMS) under the Muscle Cell Biology, Pathophysiology, and Therapeutics Training Grant (T32 AR065972). J.P. received additional funding via the Zamenhof Scholarship in Biological Chemistry and a UCLA Dissertation Year Award. B.T.N. was supported by a Philip Whitcome Pre-doctoral Fellowship from the UCLA Molecular Biology Institute.

## Author contributions

G.S., F.B., J.P., S.F.N., M.C.M., and F.G. designed research; G.S., F.B., J.P., S.M., and I.L. performed research; G.S., F.B., J.P. S.F.N., M.C.M., and F.G. contributed new reagents/analytic tools; G.S., F.B., J.P., S.M., S.F.N., M.C.M. and F.G. analyzed data; B.T.N. conducted structural modeling of the 3D-bridge ASO in complex with the exon 44 RNA; and G.S., F.B., J.P., S.F.N., M.C.M., and F.G. wrote the paper.

