## Supplemental Information for "Structural engineering of dual-targeting antisense oligonucleotides enhances exon 44 skipping for treating Duchenne muscular dystrophy"

**This PDF file includes Figures S1 to S3**

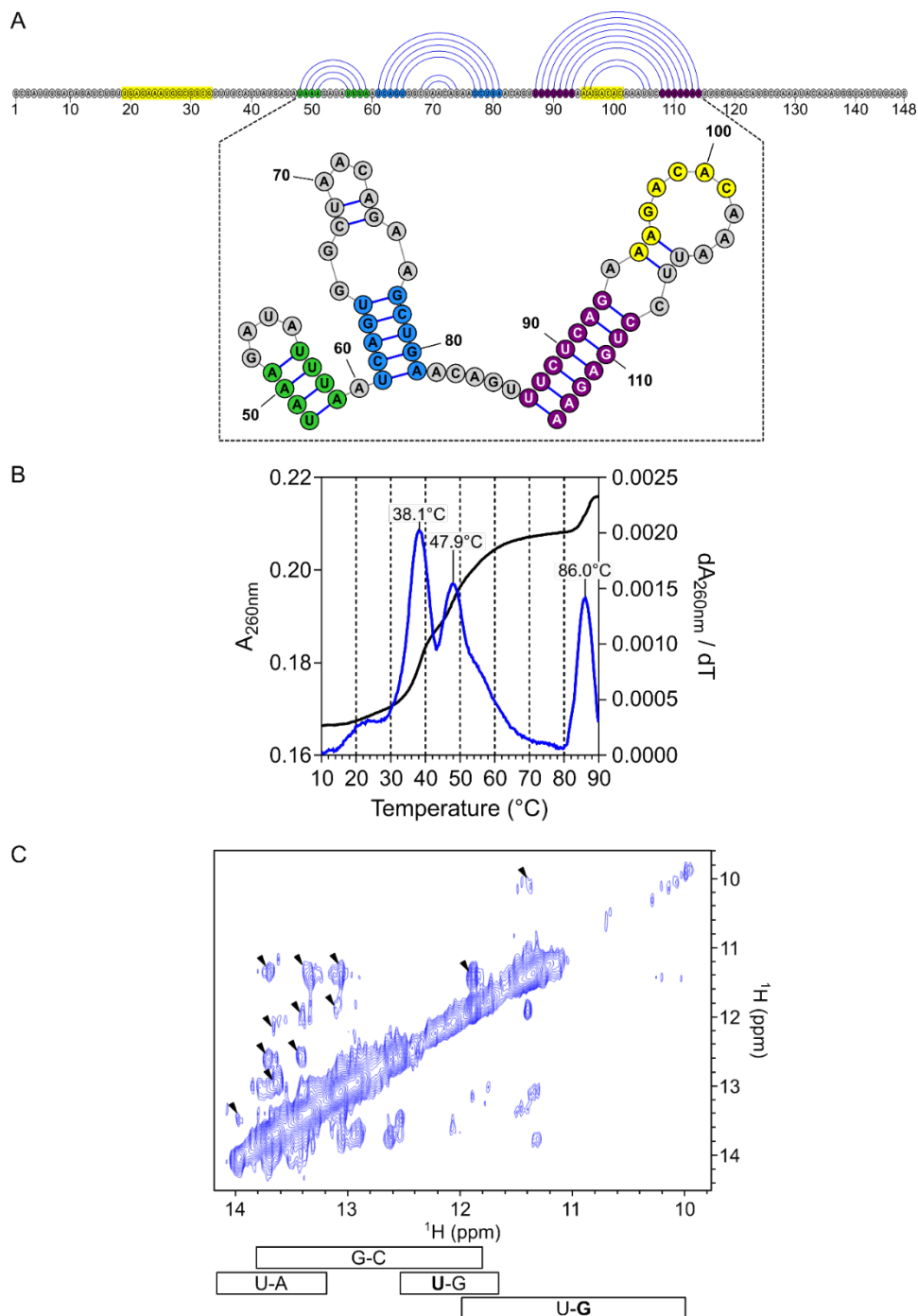

**Fig. S1.** Probing the folding of exon 44. (A) Predicted secondary structure of exon 44 in the vicinity of residues 48–114. Long-range interactions predicted between the 5' and 3' ends of the exon are not shown. Three putative stem-loop elements are colored green, blue, and purple. Regions that make Watson-Crick interactions with the 3D-bridge ASO are highlighted in yellow (residues 19–34 and 95–101). (B) UV-melting experiment with purified exon 44 RNA. The melting curve recorded at 260 nm is shown in black and the first derivative of the curve is in blue. (C) Imino proton region of the NOESY spectrum of E44<sup>19-101</sup> RNA collected at 5°C. Cross-peaks representing base pairs are indicated with arrowheads. Approximate chemical shift ranges for Watson-Crick and G-U wobble pairs are shown.

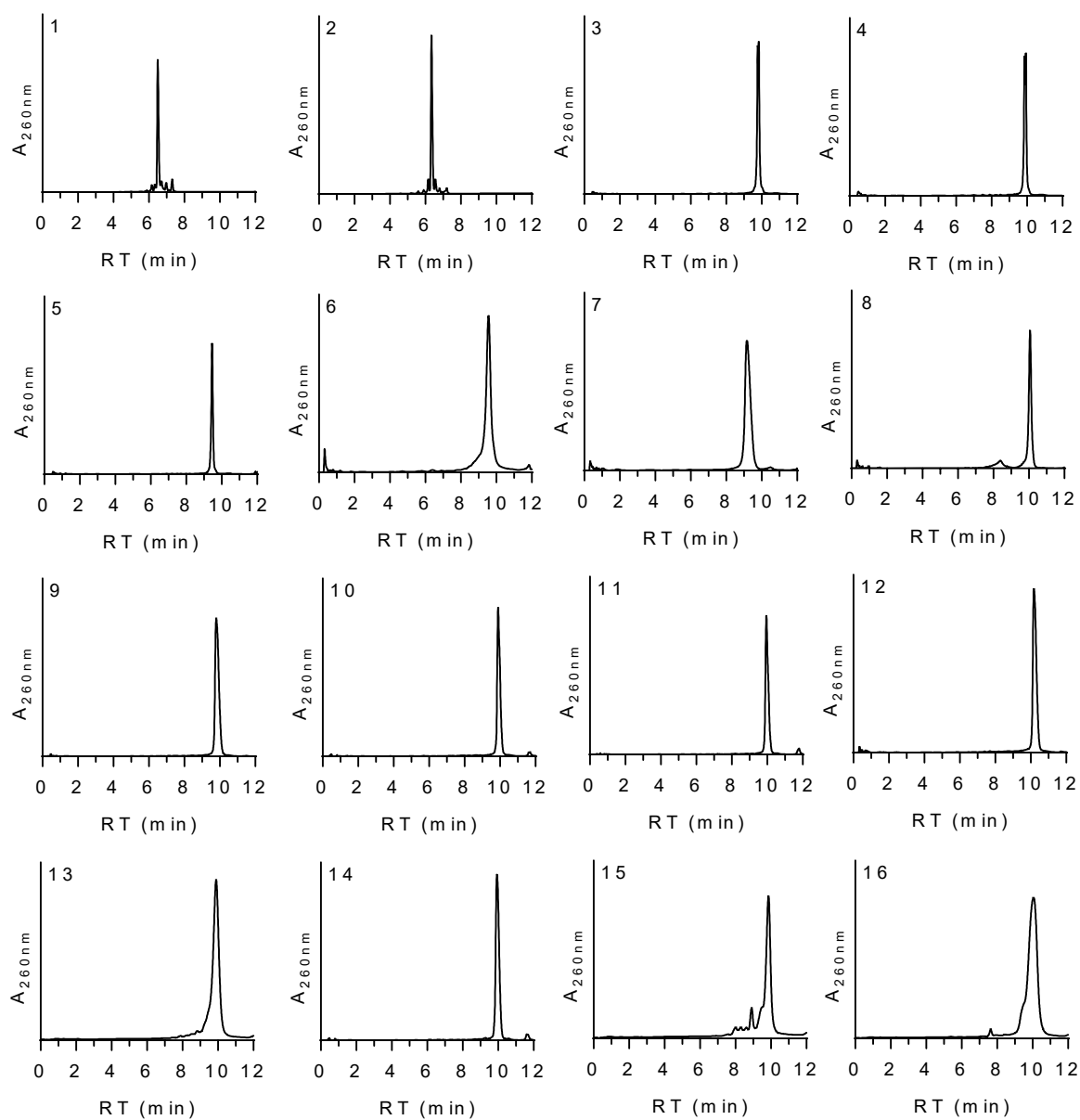

**Fig. S2.** Representative LC-MS chromatograms showing single peak profiles for the 16 oligos outlined in Table 1.

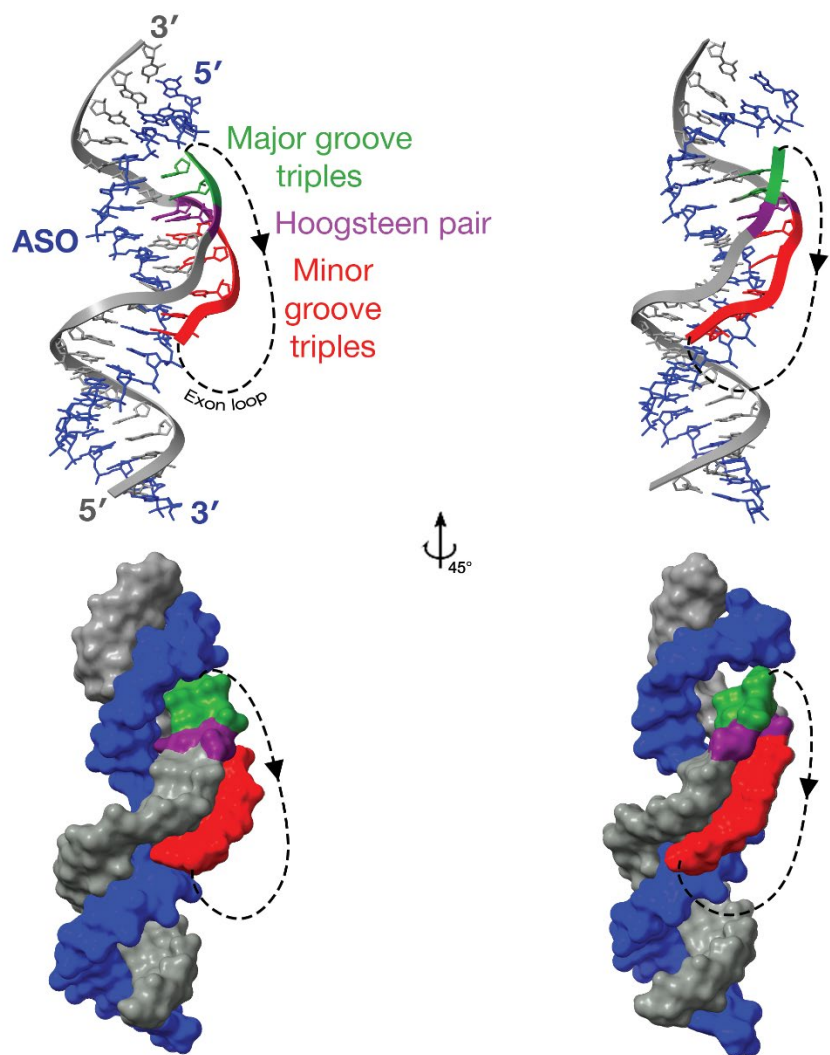

**Fig. S3.** Model of 3D-bridge-LNA/DNA bound to exon 44. Similar to Fig. 1D, the ASO strand is blue. Target RNA residues are colored according to their primary interactions, gray for Watson-Crick pairing, green for major-groove base triples, magenta for Hoogsteen base pairing, and red for minor-groove base triples. The exon loop between the two exon sites are represented by dashed lines.
